## Supplementary Materials for "Static Three-Dimensional Structures Determine Fast Dynamics Between Distal Loci Pairs in Interphase Chromosomes"

### I. DERIVATION OF DYNAMICAL PROPERTIES

We provide a brief derivation of Eq. (3) in the main text. The method used is essentially the same as in study of the dynamics of the Rouse model [1] except we use the numerically calculated $\boldsymbol{K}$ matrix and the three dimensional coordinates that are calculated using the Hi-C data as input (see Eq. (2) in the main text). We assume that the chromatin loci are subject to Brownian dynamics. Each locus experiences the same friction coefficient. With this assumption, the equations of motion of chromatin loci are given by,

$$\xi \frac{\mathrm{d}\boldsymbol{R}}{\mathrm{d}t} = \boldsymbol{K}\boldsymbol{R} + \boldsymbol{f} \tag{1}$$

where $\xi$ is the friction coefficient, $\boldsymbol{R} = (\boldsymbol{r}_1, \boldsymbol{r}_2, \cdots, \boldsymbol{r}_n)^T$ with $\boldsymbol{r}_i$ being the position of the $i^{th}$ locus, $\boldsymbol{f}$ is the Gaussian random force, and $\boldsymbol{K}$ is the connectivity matrix. By exploiting the quadratic form of the effective interaction (Eq. 2 in the main text), we perform an eigendecomposition of $\boldsymbol{K}$ as,

$$\boldsymbol{V}\boldsymbol{K}\boldsymbol{V}^T = \boldsymbol{\Lambda} \tag{2}$$

where $\boldsymbol{\Lambda}$ is the diagonal matrix with diagonal elements $\lambda_0 \geq \lambda_1 \geq \cdots \geq \lambda_{n-1}$, and $\boldsymbol{V}$ is the corresponding eigenvector. $\lambda_0 = 0$ corresponds to the zero mode (center of system). Other eigenvalues are negative. Let us define a transformation, $\boldsymbol{X} = \boldsymbol{V}\boldsymbol{R}$ where $\boldsymbol{X}$ is the normal mode. It is clear that $\boldsymbol{X}$ also follows Brownian dynamics,

$$\xi \frac{\mathrm{d}\boldsymbol{X}}{\mathrm{d}t} = \boldsymbol{\Lambda}\boldsymbol{X} + \tilde{\boldsymbol{f}}. \tag{3}$$

Because $\boldsymbol{\Lambda}$ is a diagonal matrix, the modes $\boldsymbol{X}$ are independent of each other. The independent modes follow an Ornstein-Uhlenbeck process, allowing us to write the relaxation associated with the normal modes as,

$$X_{p,\alpha}(t) \sim \mathcal{N}(X_{p,\alpha}(0)e^{-t/\tau_p}, -\frac{k_B T}{\lambda_p}(1 - e^{-2t/\tau_p})) \tag{4}$$

where $\alpha = (1, 2, 3)$ labels the three coordinates of $\boldsymbol{X}$, $\tau_p = -\xi/\lambda_p$ is the relaxation time of $p^{th}$ mode, $\mathcal{N}(\cdot)$ represents normal distribution (the first argument in $\mathcal{N}(\cdot)$ is the mean and

*
†

the second is the variance). The vector between the $i^{th}$ loci and $j^{th}$ loci is, $\boldsymbol{r}_{ij} = \boldsymbol{r}_i - \boldsymbol{r}_j$, which can be expressed using normal modes as,

$$\boldsymbol{r}_{ij} = \sum_{p=0}^{N-1} (V_{pi} - V_{pj}) \boldsymbol{X}_p \tag{5}$$

$V_{pi}$ is the matrix elements of $\boldsymbol{V}$. Since $\boldsymbol{r}_{ij}$ is a linear combination of normal modes $\boldsymbol{X}_p$, we obtain,

$$r_{ij,\alpha}(t) \sim \mathcal{N}\Big( \sum_{p=0}^{N-1} (V_{pi} - V_{pj}) X_{p,\alpha}(0) e^{-t/\tau_p}, -\sum_{p=0}^{N-1} (V_{pi} - V_{pj})^2 \frac{k_B T}{\lambda_p} (1 - e^{-2t/\tau_p}) \Big) \tag{6}$$

$$\equiv \mathcal{N}(\mu_{ij}(t), \sigma_{ij}^2(t)). \tag{7}$$

In the above equation,

$$\mu_{ij}(t) = \sum_{p=0}^{N-1} (V_{pi} - V_{pj}) X_{p,\alpha}(0) e^{-t/\tau_p}, \tag{8}$$

$$\sigma_{ij}^2(t) = -\sum_{p=0}^{N-1} (V_{pi} - V_{pj})^2 \frac{k_B T}{\lambda_p} (1 - e^{-2t/\tau_p}). \tag{9}$$

Now, we can derive an expression for $\langle \boldsymbol{r}_{ij}(t) \cdot \boldsymbol{r}_{ij}(0) \rangle$.

$$\langle \boldsymbol{r}_{ij}(t) \cdot \boldsymbol{r}_{ij}(0) \rangle = \sum_{\alpha} \langle r_{ij,\alpha}(t) r_{ij,\alpha}(0) \rangle \tag{10}$$

$$= 3 \sum_{p=0}^{N-1} (V_{pi} - V_{pj})^2 \exp(-t/\tau_p) \langle X_{p,\alpha}^2 \rangle \tag{11}$$

$$= 3 \sum_{p=0}^{N-1} (V_{pi} - V_{pj})^2 \exp(-t/\tau_p) \big( -\frac{k_B T}{\lambda_p} \big) \tag{12}$$

The two-point mean square displacement $M_2(t)$ is given by,

$$M_2(t) = 2\langle \boldsymbol{r}_{ij}^2 \rangle - 2\langle \boldsymbol{r}_{ij}(t) \cdot \boldsymbol{r}_{ij}(0) \rangle, \tag{13}$$

where $\langle \boldsymbol{r}_{ij}^2 \rangle$ is the long-time value of the distance between loci $i$ and $j$, Similarly, for single-loci diffusion $M_1(t)$, we obtain,

$$M_1(t) = 2\langle \boldsymbol{r}_i^2 \rangle - 2\langle \boldsymbol{r}_i(t) \boldsymbol{r}_i(0) \rangle \tag{14}$$

where $\langle \boldsymbol{r}_i^2 \rangle$ is the long time value of the square of the position of the $i^{th}$ locus.

### II. EXACT SIMULATION METHODS

To obtain the First Passage Time (FPT) for contacts, we perform Brownian dynamics simulations using the HIPPS-DIMES method. The equations of motion can be decoupled into a series of independent equations for the normal modes. Each normal mode follows Brownian dynamics independently of others and can be modeled as an Ornstein-Uhlenbeck (OU) process. In this study, we employ the exact simulation method, first formulated by Gillespie [2].

The one-dimensional OU process is described by the stochastic differential equation:

$$dX(t) = \theta(\mu - X(t))dt + \sigma dW(t), \tag{15}$$

where $\theta$ is the rate of mean reversion, $\mu$ is the long-term mean, $\sigma$ is the volatility, and $W(t)$ is a Wiener process.

The exact solution for this equation can be derived using the transition density function, resulting in,

$$X(t) = X(0)e^{-\theta t} + \mu(1 - e^{-\theta t}) + \sigma e^{-\theta t} \int_0^t e^{\theta s} dW(s). \tag{16}$$

The integral with respect to the Wiener process in equation given above can be expressed as a stochastic integral, and its realization can be generated using a Gaussian random variable. The final expression for simulating $X(t)$ starting from $X(0)$ is:

$$X(t) = X(0)e^{-\theta t} + \mu(1 - e^{-\theta t}) + \sqrt{\frac{\sigma^2}{2\theta}(1 - e^{-2\theta t})}Z, \tag{17}$$

where $Z$ is a standard normal random variable.

Now we can apply Eq.(17) to our model. By considering Eq.(3) in the HIPPS-DIMES model and comparing it to Eq.(15), we find that $\mu = 0$, as there is no drift term; $\theta = \Lambda/\xi$ where $\Lambda$ is the diagonal matrix of eigenvalues; and $\sigma = \sqrt{2/(\xi\beta)}$ where $\xi$ is the friction coefficient and $\beta = 1/k_B T$. Therefore, we can use Eq.(17) to evolve each normal mode with a timestep $\Delta t$, updating $X(t + \Delta t)$ from $X(t)$ using Eq.(17). Note that $\Delta t$ need not be small because Eq.(17) provides an exact update. Consequently, we can use a large timestep $\Delta t$ to speed up the simulations. At each timestep, the actual coordinates of the chromatin loci are computed from the normal modes using $\boldsymbol{R} = \boldsymbol{V}^T \boldsymbol{X}$. Without loss of generality, we set $\xi = 1$ and $\beta = 1$.

By using the numerically exact method, we can accurately simulate the dynamics of each normal mode within the HIPPS-DIMES framework. To obtain the first passage time (FPT) for contact formation, we simulate chromatin dynamics using Eq.(17). At each time step, we compute the pairwise distance between targeted pair of loci. If the distance falls below a predefined contact threshold for contact for the first time, we record the current time step as the FPT for that contact. As mentioned before, Eq.(17) is exact, thus allowing us to use a relatively large time step $\Delta t$ to accelerate the simulations without sacrificing accuracy. This approach combined with the correction method for FPT described in the section IV ensures that the mean value of FPT of contacts is computed with high precision.

### III. POLYMER SIMULATIONS

For the simulations of polymers in good and poor solvent conditions, we performed Brownian Dynamics (BD) simulations using LAMMPS [3]. Two types of chains were considered: a self-avoiding walks (polymers in a good solvent) modeled using the Weeks-Chandler-Andersen (WCA) potential [4], and a collapsed chain (polymers in a poor solvent) modeled using the Lennard-Jones (LJ) potential with $\epsilon = 1.0\, k_B T$. The chain length in both cases was taken to be $L = 64$. Simulations were conducted in the LJ unit system. Hence, all the quantities are dimensionless.

The equation of motion for the BD simulations is the overdamped Langevin equation,

$$\dot{\mathbf{r}}_i = \frac{\mathbf{F}_i}{\xi} + \sqrt{\frac{2k_B T}{\xi}}\, \boldsymbol{\eta}_i(t), \tag{18}$$

where $\mathbf{r}_i$ is the position of the $i$-th bead, $\mathbf{F}_i = -\nabla U_i$ is the systematic force derived from the potential energy, $\xi$ is the friction coefficient, and $\boldsymbol{\eta}_i(t)$ is a Gaussian random force satisfying $\langle \boldsymbol{\eta}_i(t) \rangle = 0$ and $\langle \boldsymbol{\eta}_i(t) \cdot \boldsymbol{\eta}_j(t') \rangle = \delta_{ij}\delta(t - t')$.

To model the bonds connecting two consecutive between the beads along the polymer chain are connected we use via a finite extensible nonlinear elastic (FENE) bond potential ($U_{\text{FENE}}$). In addition, beads separated by a distance $r$ interact via the WCA potential (polymer in a good solvent) or by a Lennard-Jones potential. The FENE potential is given by,

$$U_{\text{FENE}}(r) = -\frac{1}{2}KR_0^2 \ln\left(1 - \left(\frac{r}{R_0}\right)^2\right),$$

where $K = 30$, $R_0 = 1.5$, $\epsilon = 1.0$ and $\sigma = 1.0$. The specific parameters values are chosen to prevent bond crossing and ensure chain connectivity [5].

For the self-avoiding chain (SAW), we used the the WCA potential to model steric repulsion between beads:

$$U_{\text{WCA}}(r) = \begin{cases} 4\epsilon\left[\left(\frac{\sigma}{r}\right)^{12} - \left(\frac{\sigma}{r}\right)^{6} + \frac{1}{4}\right], & r \leq 2^{1/6}\sigma, \\ 0, & r > 2^{1/6}\sigma. \end{cases} \tag{19}$$

The total potential is given by $U_T = U_{\text{FENE}} + U_{\text{WCA}}$.

For the collapsed chain, the LJ potential with $\epsilon = 1.0\, k_BT$ captured both the attractive and repulsive interactions:

$$U_{\text{LJ}}(r) = 4\epsilon\left[\left(\frac{\sigma}{r}\right)^{12} - \left(\frac{\sigma}{r}\right)^{6}\right]. \tag{20}$$

As in good solvents, the total potential is, $U_T = U_{\text{FENE}} + U_{\text{LJ}}$. The simulation parameters included a temperature of $k_BT = 1.0$, interaction strength $\epsilon = 1.0$ (for WCA/LJ potential), bead diameter $\sigma = 1.0$, friction coefficient $\xi = 1.0$, time step $\Delta t = 10^{-4}$, and a total simulation time of $10^9$ time steps. The time step was chosen to balance numerical stability and computational efficiency.

The simulations of self-avoiding and collapsed polymer chains are used to validate the theoretical results of our model. To validate the theoretical predictions, we computed the dynamic properties $M_1(t)$ and $M_2(t)$ from the simulation trajectories of the self-avoiding and collapsed polymer chains. First, the average distance matrix was calculated from the trajectories. This average distance matrix was then fed into the HIPPS-DIMES method to obtain the connectivity matrix $\boldsymbol{K}$. Using this connectivity matrix, we used Eqs. (14) and (13) to calculate $M_1(t)$ and $M_2(t)$ theoretically. The theoretical results were compared with the corresponding quantities directly computed from the simulation trajectories. The comparisons, shown in SI Figs. 4 and 5, demonstrate excellent agreement between the simulations and the theory.

### IV. ESTIMATING THE MEAN FIRST PASSAGE TIME (MFPT) FROM FINITE TIME SIMULATIONS

Like in most if not all simulations, the First Passage Time (FPT) for contact is calculated using a maximum number of time steps $T$ due to computational limitations. In many of the trajectories, contacts between various loci may not form on the time, $T$. In such instances, we only know that the FPT exceeds $T$, but the exact values are unknown. Simply averaging the observed FPTs over multiple trajectories or by increasing $T$ does not guarantee convergence to the true answer because these strategies ignore the contributions from the events that occur on time scales that greatly exceed $T$. These issues are well-recognized in statistics and are referred to as the problem of *incomplete or censored data*, where some observations are only partially known. To address this bias and obtain an accurate estimate of the true mean FPT, we utilize Maximum Likelihood Estimation (MLE) for incomplete data.

**Method for Estimating the True Mean FPT (MFPT)**: Assuming that the distribution of FPTs follows an exponential distribution, the MLE estimate of the true mean FPT $\langle\tau\rangle$ can be calculated using the following formula [6]:

$$\langle\tau\rangle = \frac{\sum_{i=1}^{n} \tau_i + n_{\mathrm{E}} \cdot T}{n}, \tag{21}$$

where $\tau_i$s are the observed FPTs that are less than $T$, $n$ is the number of events that are less than $T$ (i.e., the number of simulations where the FPT was observed within the simulation time), and $n_{\mathrm{E}}$ is the number of simulations in which the FPT exceeds $T$ ( the instances where the first passage did not occur within the simulation time). The total number of simulations, $N = n + n_{\mathrm{E}}$. This formula given above adjusts the mean estimate by incorporating both the observed FPTs and the number of instances that exceed the maximum simulation time $T$. Note that if $n_{\mathrm{E}} = 0$ then the correction term is not needed. By accounting for these incomplete observations, the MLE method provides a more accurate estimate of the true mean FPT.

**Validating Eq. 21 using Synthetic Data**: To illustrate the effectiveness of this Eq.(21), we conducted a synthetic simulation where the true mean FPT is known. Let $\tau$ be a random variable that is exponentially distributed with a true mean $\bar{\tau} = 100$. We generate datasets by sampling from this distribution but limit the maximum value to $T = 80$ that is less than the true mean value. Any sampled value exceeding 80 is only recorded as "greater than 80.”

Supplementary Fig. 1 illustrates the results of these simulations. In panel (a), we present the empirical distributions of a random variable $\tau$ drawn from an exponential distribution with a true mean $\bar{\tau} = 100$, but with observations limited to $\tau \leq 80$ (reflecting a maximum simulation time $T = 80$). This means that any value of $\tau$ exceeding 80 is not observed. The left plot in Fig. 1(a) corresponds to a sample size of $N = 100$. The empirical distribution is truncated at $\tau = 80$ which means there are instances where values $> 80$ are not sampled. The direct calculation (neglecting the second term in Eq.21) of the mean from the observed data yields $\bar{\tau} = 33.8$, which significantly underestimates the true mean due to the unobserved values of $\tau$ that exceed $T$. By applying the MLE method using Eq. (21) with known $n$ and $n_{\mathrm{E}}$ ($N = n + n_{\mathrm{E}}$) for incomplete data, we obtain an adjusted mean estimate $\bar{\tau}_{\mathrm{MLE}} = 87.13$, which is much closer to the true mean of 100.

The right plot in Fig. 1(a) shows the results for a larger sample size of $N = 10,000$. As before, the empirical distribution similarly truncates at $\tau = 80$. The naive mean estimate is $\bar{\tau} = 34.72$, while the MLE-adjusted mean is $\bar{\tau}_{\mathrm{MLE}} = 100.65$, closely matching the true mean. The larger sample size (10,000 versus 100) improves the accuracy of the MLE estimate, demonstrating its robustness even with incomplete data. This exercise hows that increasing the sample size alone is insufficient to obtain results that are close to the exact value. No matter how large $N$ is, one must use the correction in Eq. (21) to obtain an answer that is close to the expected result.

In Fig. 1(b), we show the estimates of the mean as a function of the maximum observed value $T$ (the threshold beyond which exact values of $\tau$ are unknown), ranging from $T = 10$ ($< \bar{\tau}$) to $T = 500$ ($> \bar{\tau}$). The blue data points represent the mean estimates calculated directly from the observed data without corrections given in Eq. 21. These estimates increase monotonically with $T$ but only approach the true mean when $T$ exceeds 300, indicating that very long simulation times would be required to accurately estimate the mean without the MLE correction. In contrast, the green data points represent the MLE-adjusted mean estimates as a function of $T$. Even for small values of $T$, the MLE method provides mean estimates close to the true mean of 100. This demonstrates that the MLE method effectively corrects for the bias introduced by restricting the observation time, allowing accurate estimation of the true mean even when $T$ is small.

**Application to Simulation Data**: Building on the effectiveness of the Maximum Like-

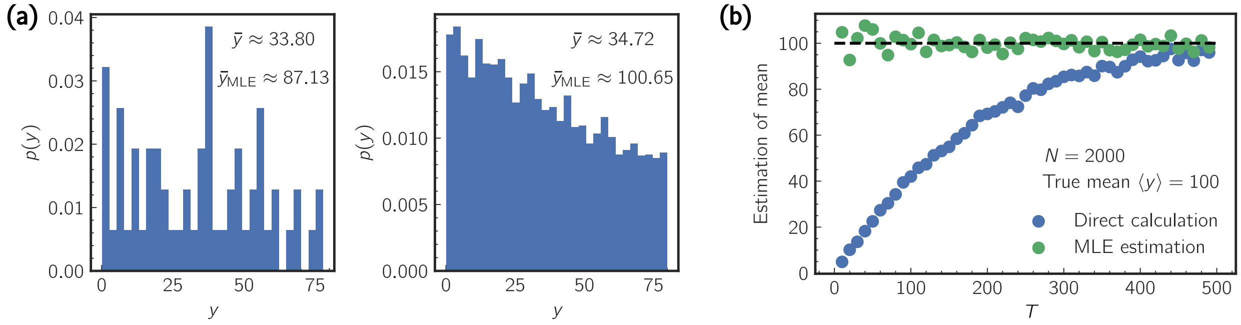


FIG. 1. Efficacy of the MLE method (Eq. (21)) using synthetic data with known true mean $\langle y \rangle = 100$. (a) Empirical distributions of $y$ drawn from an exponential distribution that is limited to $y \leq 80$ (censoring at $b = 80$). Left: Sample size $N = 100$; the naive mean is $\bar{y} = 33.8$, and the MLE-adjusted mean is $\bar{y}_{\mathrm{MLE}} = 87.1$. Right: Sample size $N = 10,000$; the naive mean is $\bar{y} = 34.7$, and the MLE-adjusted mean is $\bar{y}_{\mathrm{MLE}} = 100.7$. (b) Estimates of the mean as a function of the threshold $b$. Blue points: Naive mean estimates, which increase with $b$ but only approach the true mean when $b$ is large. Green points: MLE-adjusted mean estimates, which remain close to the true mean even for small $T$ values.

lihood Estimation (MLE) method demonstrated with synthetic data, we used this approach to actual simulation data of first passage times (FPTs) for contact between pairs of loci. To generate the simulation data, we employed the exact simulation method described earlier, using Eq.(17) to model the dynamics of each normal mode in the HIPPS-DIMES model. Starting from an initial configuration at equilibrium, we evolved the system over time using a time step $\Delta t = 0.1$ (with $\xi = 1$ and $\beta = 1$) for a total number of time steps $T$. At each time step, we updated the normal modes according to Eq.(17) and computed the actual coordinates of the chromatin loci using $\boldsymbol{R} = \boldsymbol{V}^T \boldsymbol{X}$, which allows us to calculate the pairwise distances between all the loci. If the distance between any pair of loci fell below the predefined contact threshold, $r_c = 1$, for the first time, we recorded the current time as the FPT for that contact. If no contacts occurred up to the end of the simulation at time $T$, we recorded the FPT as being greater than $T$. By repeating this simulation multiple times, we collected samples of observed FPTs for various pairs of loci, including records of cases where no contact occurred during the simulation. Here, we show that the MLE method improves the estimation of the mean first passage time (MFPT) in our simulations.

Figure 2(a) shows the empirical survival probability $S(\tau_c)$ of the FPTs, $\tau_c$, for contact

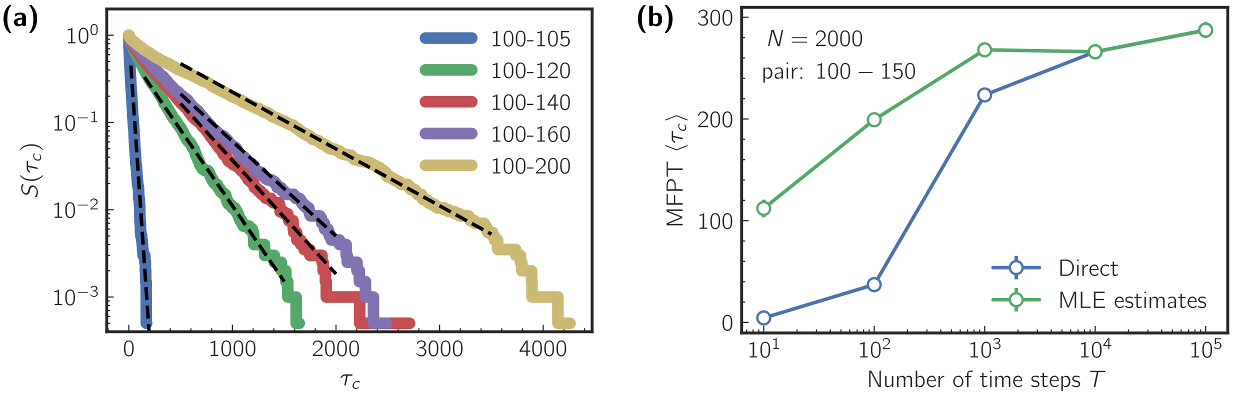


FIG. 2. Application of the MLE method to simulation data for estimating the mean first passage time (FPT) between loci. (a) Survival probability of FPTs for contact between several pairs of loci $S(\tau_c)$, plotted on a semi-logarithmic scale. The straight lines indicate an exponential distribution of FPTs. Dashed lines are guides for eye. (b) Estimates of the mean FPT $\langle\tau_c\rangle$ as a function of the maximum number of time steps $T$. Blue points: Direct mean estimates, which saturate and underestimate the true mean at large $T$. Green points: MLE-adjusted mean estimates, which provide accurate estimates even at smaller $T$.

between several pairs of loci, plotted on a semi-logarithmic scale (logarithm of the probability density function on the y-axis versus linear FPT on the x-axis). The linearity of these plots shows that the FPTs closely follow an exponential distribution, supporting the assumption underlying the MLE method. Figure 2(b) illustrates the estimates of the mean FPT, $\langle\tau_c\rangle$, as a function of the maximum number of time steps $T$ for a specific loci pair, (100 - 150). Two sets of estimates are shown: Direct estimates (blue data points), calculated by averaging the observed FPTs within the simulation time $T$, and MLE Estimates (green data points), calculated using Eq. (21), which account for FPTs exceeding $T$. The direct estimates saturate at large $T$. Although the true mean is not known, the saturation at large $T$ suggests that the true mean is approximately 260. At small $T$, the direct estimates significantly underestimate the true mean due to the exclusion of longer FPTs. In contrast, the MLE estimates improve the accuracy of the mean FPT, providing values closer to the true mean even at smaller $T$.

The comparison highlights that the MLE method effectively corrects the bias inherent in the finite simulation times, leading to more accurate estimation of $\langle\tau_c\rangle$ without the need for excessively long simulations.

**Application to Experimental Trajectories:** The Maximum Likelihood Estimation (MLE) method described earlier can be applied to experimental trajectory data, where the maximum observation time $T$ is not uniform across all the trajectories. In experiments the trajectories often vary in length due to practical constraints, with some being shorter than the others. This non-uniformity means that each trajectory may have its own maximum observation time $T_i$. To account for this variation, Eq. (21) can be modified to incorporate individual observation times for each trajectory. Specifically, the MLE formula is adjusted to sum over the observed FPTs and the maximum observation times for trajectories where the FPT exceeds their respective $T_i$. The modified formula is given by,

$$\langle \tau \rangle = \frac{\sum_{i=1}^{n} \tau_i + \sum_j T_j}{n}, \tag{22}$$

where $T_j$ is the length of trajectory $j$ where the first passage event does not occur. By using the above equation, we can then estimate the mean FPT from experimental data, effectively correcting for biases introduced by incomplete observations due to finite and non-uniform observation times.

We applied the modified MLE method to estimate the mean FPT from experimental trajectory data [7]. The estimated mean FPTs were then compared to the predictions from our theoretical model presented in Fig. 5b in the main text. The close agreement between the experimental estimates and the model predictions validates the applicability of the MLE method to experimental data and further supports the robustness of our theory.

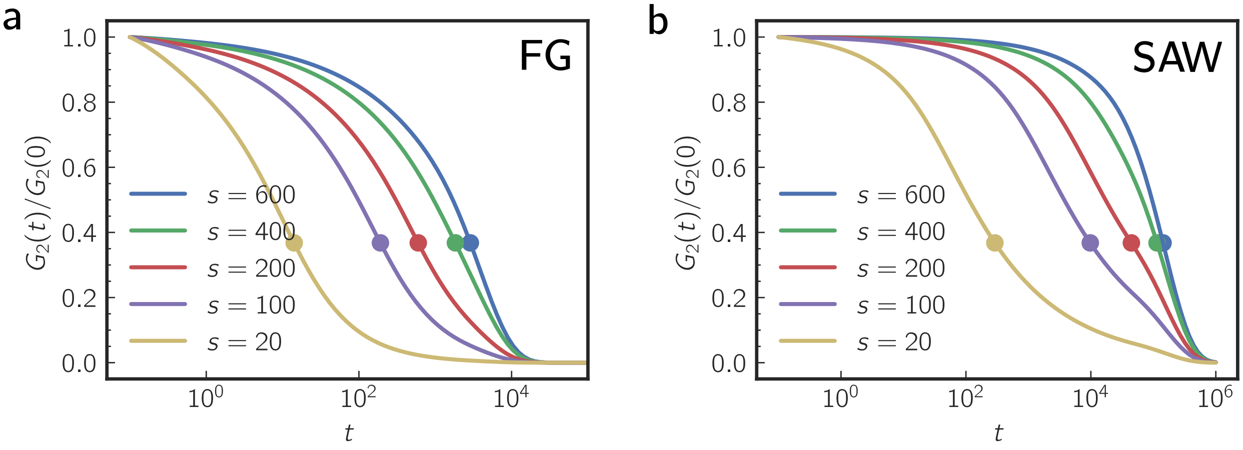


FIG. 3. (a) Normalized two-point relaxation time, $G_2(t)/G_2(0)$, for different sub-chains of length $s$. The results are for the fractal globule with chain length $N = 1,000$. Symbols indicate the location of the relaxation time $\tau$, defined as $G_2(\tau)/G_2(0) = 1/e$. (b) Same as (a) but for the self-avoiding chain model.

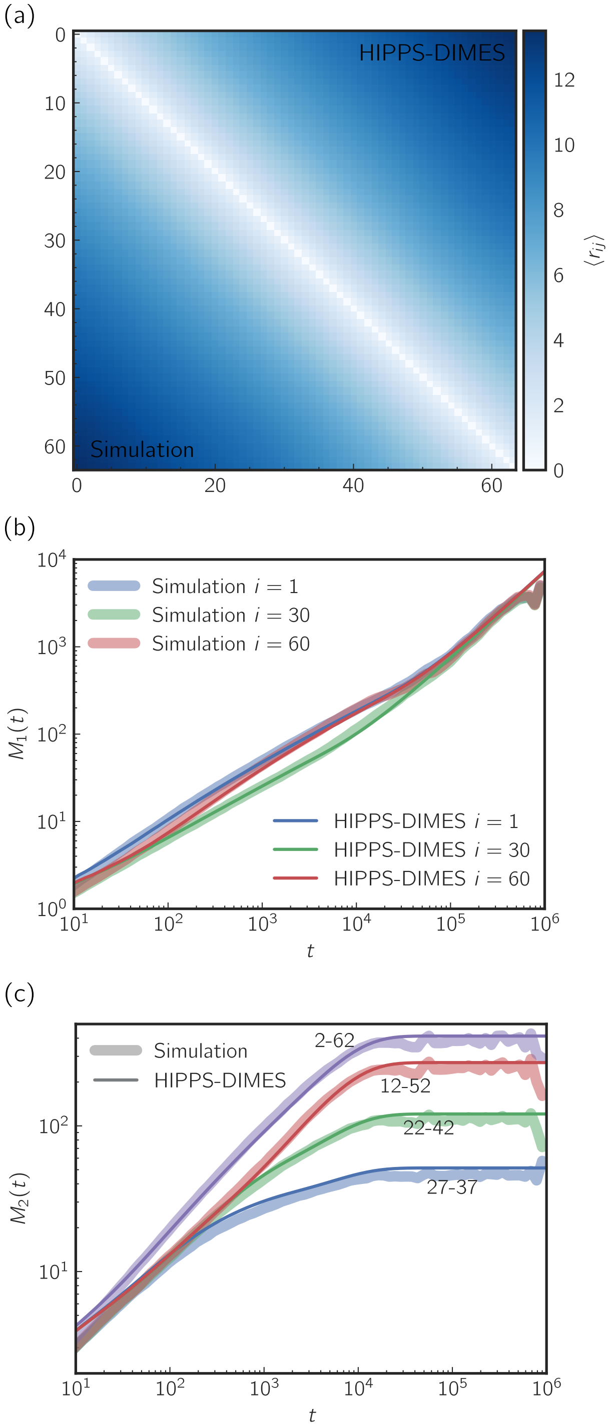


FIG. 4. Comparison between the simulation and HIPPS-DIMES for Self-avoiding Chain (SAW), a polymer in a good solvent. Chain length is $L = 64$. (a) Comparison of the mean distance map between simulation and the model. (b) Comparison of Mean Square Displacement of single monomer $M_1(t)$ between simulation and the theory, where $i$ is the index of the monomer. The agreement between theory and simulations is excellent. (c) Time-dependent changes in the two-point Mean Square Displacement $M_2(t)$ between simulations and the theory. The numbers indicate the indices of the monomer pair.

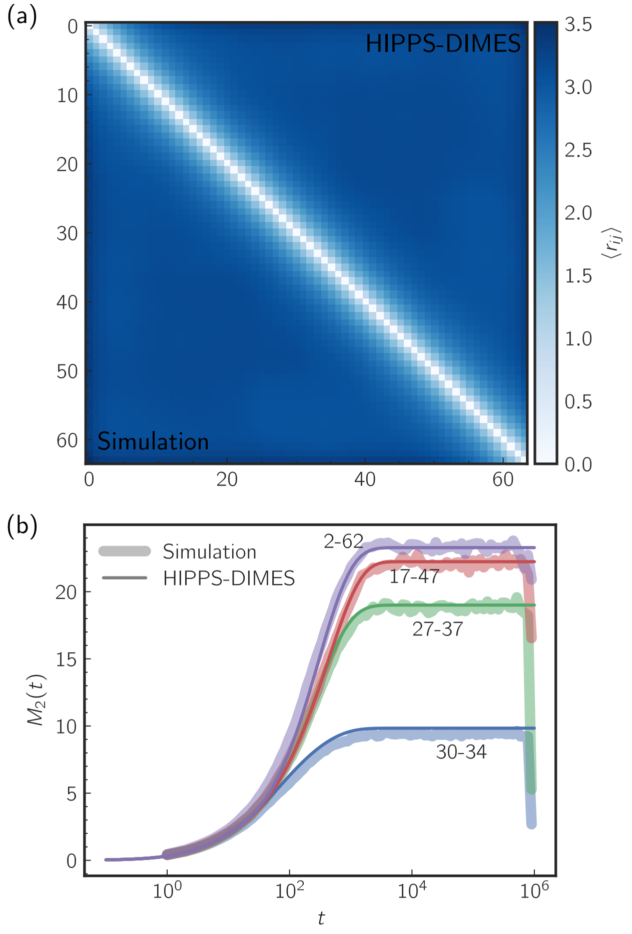


FIG. 5. Comparison between the simulation and HIPPS-DIMES for collapsed chain (polymer in a bad solvent). Chain length is $L = 64$. The monomer-monomer interactions are modeled as Lennard-Jones potential with interaction strength $\epsilon = 1.0k_BT$ (see Eq. 20). (a) The mean distance map calculated using theory is in excellent agreement with simulations. (b) Comparison of two-point Mean Square Displacement $M_2(t)$ between theoretical predictions and simulations. The numbers correspond to the indices of the monomer pair.

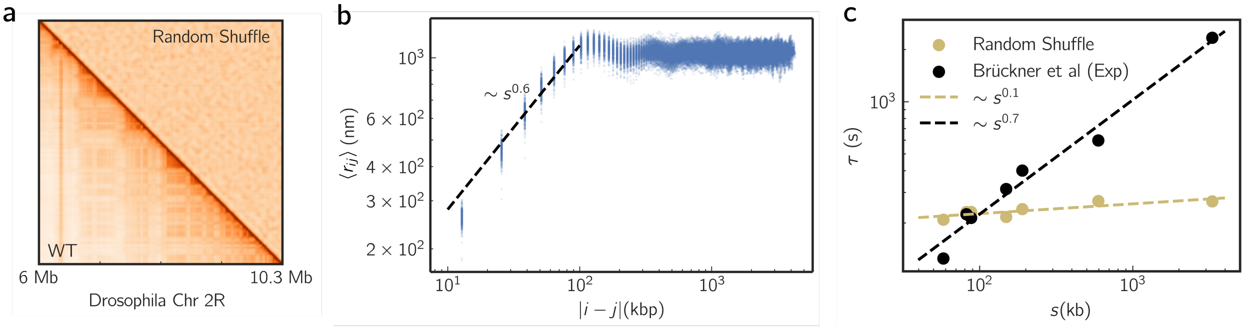


FIG. 6. (a) Side-by-side comparison of the contact map between the wild type (WT) Drosophila Chromosome 2R and a randomly shuffled contact map. We shuffled the values of the off-diagonal elements that are at least two positions away from the main diagonal (i.e., elements where $|i-j| \geq 2$), while keeping the matrix symmetric. Specifically, we collected these values, randomly permuted them, and reassigned them symmetrically to their original positions. This method preserved immediate neighbor relationships while randomizing longer-range connections in the matrix. (b) Mean pairwise distance $\langle r_{ij} \rangle$ as a function of genomic distances $|i-j|$. (c) Relaxation time $\tau$, versus the corresponding genomic distances $s$. The dependence of $\tau(s)$ on $s$ for the shuffled sequence is dramatically different from the WT.

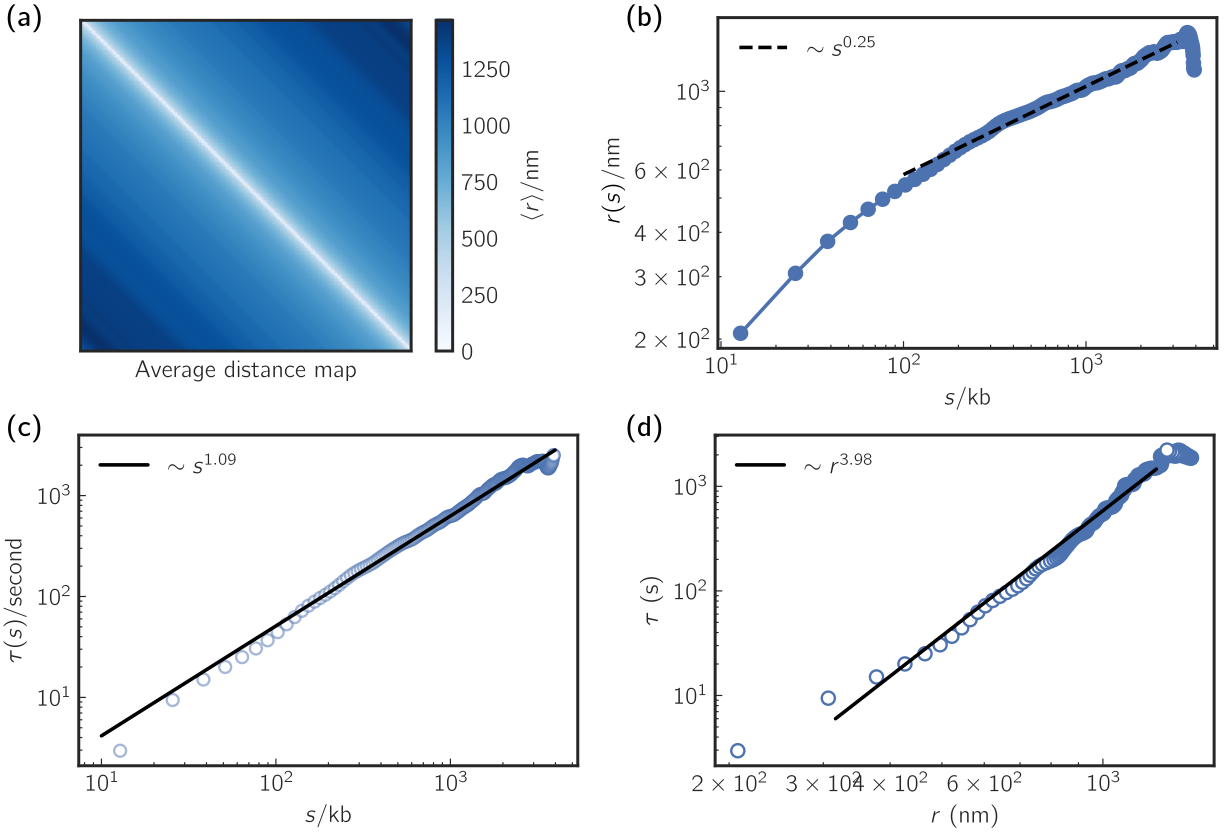


FIG. 7. (a) Average distance map for an effective homopolymer model. Each entry is calculated as $r(s=|j-i|)=\sum_{|j-i|=s}\langle r_{ij}\rangle/(N-s)$ where $N$ is the total number of locus and $s$ is the genomic distance. (b) Scaling of $r(s)$ as a function of $s$. (c) Scaling of relaxation time $\tau$ as a function of $s$. (d) Scaling of relaxation time $\tau$ as a function of mean distance $r$.

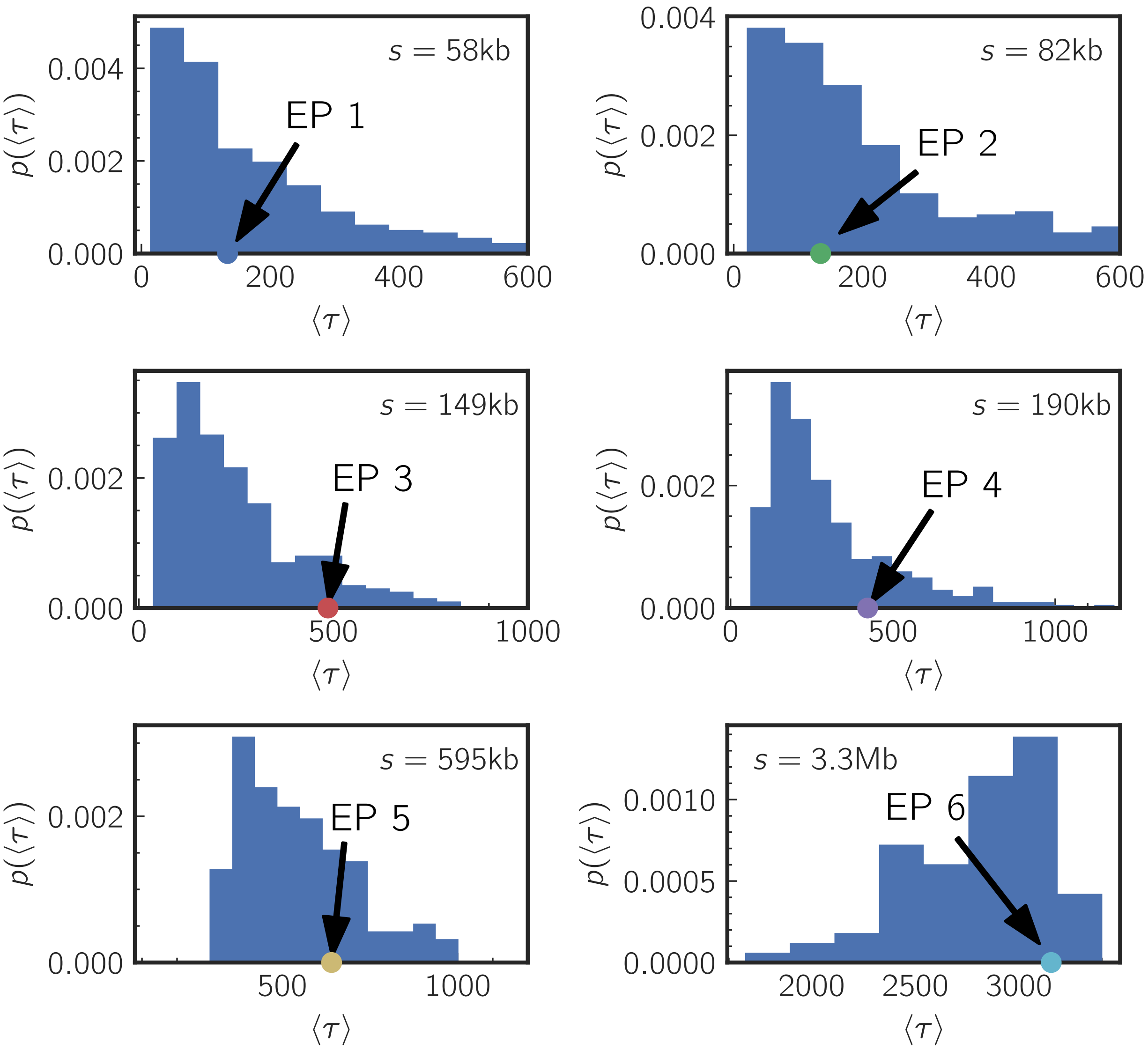


FIG. 8. Comparison of the distribution of relaxation times ($\langle\tau\rangle$) for all locus pairs with genomic distances matching those of enhancer-promoter ($\mathrm{EP}_i$ $i = 1, 2 \cdots 6$) pairs probed in experiments. Each plot shows the probability distribution ($p(\langle\tau\rangle)$) for loci at equivalent genomic separations ($s$), with the mean relaxation time for each specific enhancer-promoter pair indicated as a distinct marker.

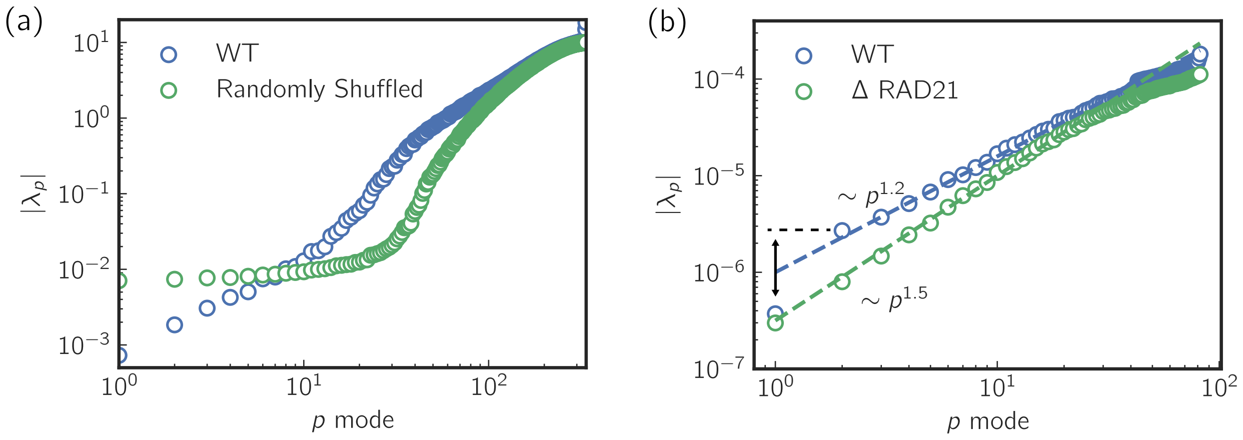


FIG. 9. (a) Comparison of the eigenvalues $|\lambda_p|$ as a function of $p$ ($p$ labels the normal mode) between Wild-type (WT) and the randomly shuffled sequence. (b) Comparison of the eigenvalues $|\lambda_p|$ as a function of $p$ ($p$ labels the normal mode) between WT and the cohesin-depleted cells ($\Delta$RAD21). The arrow marks the gap between $p = 1$ mode and $p = 2$ mode for WT cells.

[6] J. P. Klein and M. L. Moeschberger, *Survival analysis: techniques for censored and truncated data* (Springer Science & Business Media, 2006).

[7] D. B. Brückner, H. Chen, L. Barinov, B. Zoller, and T. Gregor, Stochastic motion and transcriptional dynamics of pairs of distal dna loci on a compacted chromosome, Science **380**, 1357–1362 (2023).
